# Transgenerational Effects of Early-Life Stress on Anxiety in Zebrafish (*Danio rerio*)

**DOI:** 10.1101/2022.11.22.517541

**Authors:** Barbara D. Fontana, Nancy Alnassar, Matthew O. Parker

## Abstract

Early-life adversity impacts on anxiety-related behaviors in adulthood. The effects of such adversity not only affects the animal itself, but can be passed on transgenerationally. Pervasive effects of experimentally-induced early-life stress (ELS) have been documented in adult zebrafish but it is not clear if this can be passed on via the germline. Here, we investigated the effects of ELS across three generations, by analyzing the responses of adult animals exposed to ELS in two different anxiety-related tasks, as well as in social behavior, memory, and cognition. Animals exposed to ELS (at 7 days-post-fertilization) showed a marked attenuation of specific anxiety-related behaviors (F_0_) when adults, and these alterations were maintained across two subsequent generations (F_1_ and F_2_). These findings suggest that zebrafish may be a useful model organism to study the transgenerational effects of ELS, and how this pertains to (for example) neuropsychiatric disorders. In addition, our data may naturally provoke questions regarding consideration of the environment of laboratory-housed zebrafish at early developmental stages. In particular, more work may be necessary to determine how different environmental stressors could affect data variability across laboratories.

**Graphical Abstract:** Summary of the ELS effects in zebrafish anxiety-like behavior across multiple generations.

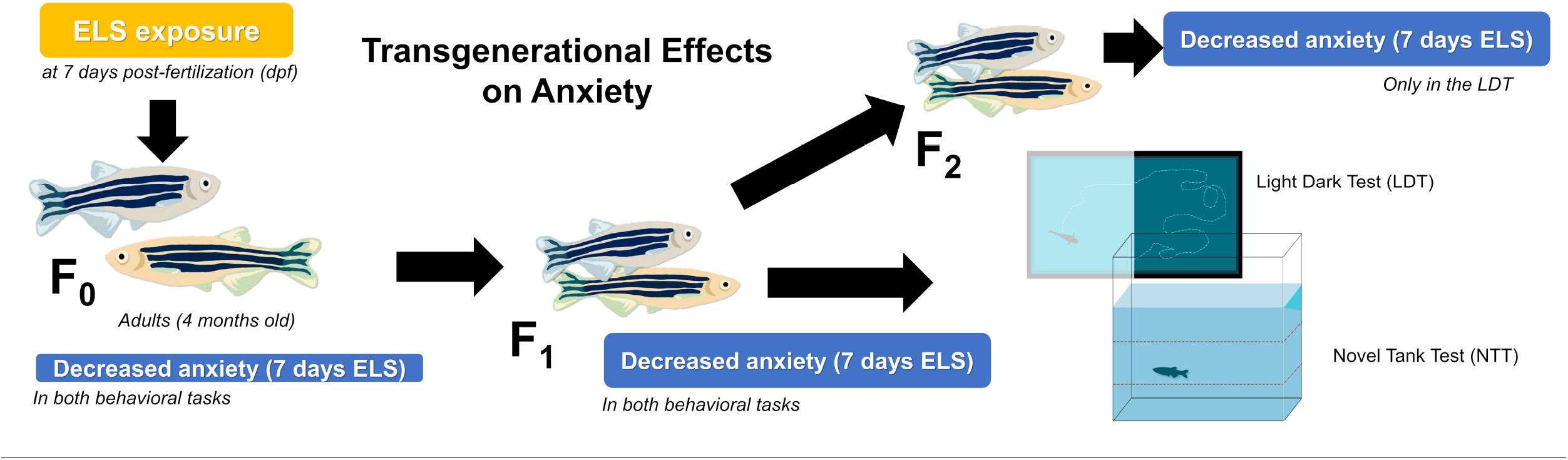

## Main

Behavioral plasticity occurs across the lifespan, and adult traits are shaped by experience. Exposure to stress or adversity during early development, for example, causes behavioral differences (compared to non-exposed matched controls) which persist throughout the lifespan, and likely reflect an adaptive response to potentially challenging ecologies (Nettle & Bateson, 2015). This is particularly pertinent in intensively managed laboratory zebrafish, where there is variability in husbandry-related early-life adversity (*e.g*., transport, changes in light, pH, water volume, and human handling, etc.)(Aleström *et al*., 2020). Behavioral and physiological adaptations that result from such early-life adversity can not only have a long-lasting impact on the individual (*i.e*., leading to variability in adult traits; (Fontana *et al*., 2021a; Fontana *et al*., 2021b)) but these effects can be found across generations in several species (Cowan *et al*., 2016). In turn, the resulting individual variability could also lead to unreliable results both within and between laboratories.

Early-life stress (ELS) can trigger positive (*e.g*. build resilience) or negative (*e.g*. risk factor for psychiatric disorders) changes which persist into adulthood (Repetti *et al*., 2002; Turner & Lloyd, 2003; Anda *et al*., 2006; Heim *et al*., 2008; Russo *et al*., 2012; Sapolsky, 2015). Behavioral and physiological changes caused by ELS can not only have long-lasting impact on the exposed individual, and may be passed to future generations (van Steenwyk & Mansuy, 2021). Complex interactions between gene expression and regulation, and the environment, are responsible for shaping behavior of organisms, and epigenetic changes during adulthood may be passed from parents to offspring (Flavell & Greenberg, 2008; Lahiri *et al*., 2016). For example, received maternal care plays a role in the subsequent maternal care given by female offspring, and this correlates with differential expression of the oxytocin receptor, suggesting a mechanism for transgenerational transmission of individual differences in stress-reactivity (Meaney, 2001). Recently, we demonstrated that exposure to chronic, mild, unpredictable ELS causes a robust decrease in anxiety-related phenotypes in adult zebrafish and altered stress-reactivity in exposed individuals (Fontana *et al*., 2021a; Fontana *et al*., 2021b). The extent to which the effects of ELS can be passed down through subsequent generations through the germline, however, has yet to be determined in zebrafish.

Here, to establish if ELS-induced adaptations in behavioral traits can be subject to germline-dependent transmission, we investigated the impacts of ELS across three generations (F_0_, F_1_ and F_2_). First, we exposed 7dpf zebrafish (F_0_) to 7 days of chronic unpredictable mild early life stress (CUELS). These fish were screened at 4 months of age for several behavioral endpoints. We then in-crossed F_0_ fish to produce F_1_, which were again grown to 4 months, and tested behaviorally. Finally, we in-crossed F_1_ to produce F_2_, and completed behavioral phenotyping at 4 months. The entire experiment was replicated in two independent batches to ensure reliability.

## Methods

Data and methods concerning the F_0_ animals has been published previously (Fontana et al 2021b), and is included here for reference.

### Animals husbandry and data reliability

AB wild-type zebrafish were bred in-house from multiple tanks (3 tanks). After egg collection, embryos were incubated in a petri dish (100 x 15 mm) until 7 dpf, then transferred to a small plastic container (15×8×2 cm length x height x water depth). At 14 dpf fish were transferred to a re-circulating (Aquaneering, USA) system in groups of 40 animals per 1.4L. Juveniles (>30dpf) were then kept in the re-circulating system in groups of 10 animals per 2.8 L until 4 months. During all life-stages animals were kept on a 14/10-hour light/dark cycle (lights on at 9:00 a.m.; pH 8.4; 28 °C (±1 °C)). Enrichment was provided at all life stages, including larvae. In larvae/juveniles this took the form of laminated images of pebbles under the tank. In older animals, several enrichment devices were placed in each tank, including laminated pebbles under the tank, and plastic plants and ‘hides’ (a hollow triangular prism on the floor of the tank) in the tanks. Animals were fed three times a day according to age following the same procedures described in Fontana et al. (2021). Behavioral experiments were performed between 10:00 and 16:00 h.

To ensure reliability, we used two independent batches for breeding and behavioral testing (*n* = 16 per group in each batch). F_0_ were reared to 4 months old and tested for social cohesion (shoal test: *n* = 8 per shoal; 1:1 sex ratio) and anxiety (novel tank diving /light-dark test: *n* = 16/group/task). Finally, F_0_ were tested for working memory and behavioral flexibility (free movement pattern [FMP] Y-maze/Pavlovian fear conditioning task: *n* = 16/group/task). A randomly selected group of F_0_ fish that were not behaviorally tested were used for breeding of F_1_; similarly, a group of F_1_ fish were just used for breeding and not tested behaviorally (*n=* 20 breeders per generation). All behavioral testing was carried out in a fully randomized order, choosing fish at random from one of three housing tanks for testing per batch. Once all data were collected and screened for extreme outliers (e.g., fish freezing and returning values of ‘0’ for behavioral parameters indicating non-engagement), the data was analyzed in full. Only one animal was excluded from F_1_ generation, group 0 days of stress exposure (see below) for the novel tank test (NTT; see below). All experiments were carried out following approval from the University of Portsmouth Animal Welfare and Ethical Review Board, and under license from the UK Home Office (Animals (Scientific Procedures) Act, 1986) [PPL: P9D87106F].

### 2.2. Chronic unpredictable early-life stress (CUELS)

The CUELS protocol comprised random exposure of two pseudo-randomly selected stressors per day at a psudo-randomly selected time. F_0_ were exposed to the CUELS protocol for 7 or 14 days depending on the group, starting when animals reached 7 dpf. The stressors consisted of social isolation (individually transfer animal to well in a white 96-well plate for 45 minutes); overcrowding (transfer 40 animals to a well in a 12-well plate for 45 minutes); light/dark cycle (light/dark cycle changes for 60 minute); water change (change animal to a new tank with new water 3 times); water cooling (transfer animal tanks to an incubator until the water temperature reaches 23 °C for 30 minutes); water heating (transfer to an incubator until the water reaches 33 °C for 30 minutes); mechanical stirring (stir the water for 5 minutes using a Pasteur pipette); immersion in shallow water (transfer animal to tanks with water removed such that the body is exposed, and leave in this condition for 2 minutes).

### 2.3. Shoaling test

Fish (4-fish per shoal; *n =* 8 shoals with 1: 1 sex ratio) were simultaneously placed in the test tank (25×15×10 cm length x height x width) and group behavior was analyzed for 5 minutes (Green *et al*., 2012; Schmidel *et al*., 2014). The Zantiks AD unit (Zantiks Ltd., Cambridge, UK) was used to record the fish behavior and videos were exported to Image J 1.49 software to assess shoaling using screenshots taken every 15 s during the 5-min trials (20 screenshots *per* trial) (Green *et al*., 2012; Schmidel *et al*., 2014). Screenshots were calibrated proportionally to the size of the tank to allow the quantification of total inter-fish distance (cm) and shoal area (cm^2^).

### 2.4. Novel tank diving test

The novel tank test is used to measure exploration, locomotion, boldness and anxiety-like behavior (Egan *et al*., 2009). Animals (*n =* 48) were placed individually in a novel tank (20 cm length x 17.5 cm height x 5 cm width) containing 1 L of aquarium water (10 cm water depth). Behavioral activity was analyzed using the Zantiks AD system (Zantiks Ltd., Cambridge, UK) for 6 minutes (Egan *et al*., 2009; Parker *et al*., 2012). The tank was separated, virtually, from bottom to top, into three zones (bottom, middle and top) to provide a detailed evaluation of vertical activity. We also measured total distance traveled (mm), number of entries and time spent in top zone (s). Finally, we measured habituation to the novel environment by considering the number of entries to, and time spent in, the top zone, across the 6-min.

### 2.5. Light-dark preference task

The light-dark preference task is a well-characterized test that aims to analyze anxiety-like behavior (Gerlai *et al*., 2000). The light-dark preference task was performed in commercially available test arena (Zantiks, Cambridge, UK), comprising a black tank (20 cm length x 15 cm height x 15 cm width) divided into two equally sized partitions where half of the tank area contained a bright white light, and the other area was covered with a black partition to avoid light exposure. Animals (*n=* 48) were place individually into the behavioral apparatus and its activity analyzed using Zantiks AD tracking systems (Zantiks Ltd., Cambridge, UK) for 6-min to determine the time spent in dark area (Blaser & Rosemberg, 2012).

### 2.6. Y-maze test

The FMP Y-maze is a validated and translationally relevant working memory task (Cleal *et al*., 2021). The FMP Y-maze apparatus is a Y-shaped insert with three identical arms (5 cm x 2 cm L x W; 120° angle between arms) with no explicit intra-maze cues, containing 3L of aquarium water. The analysis treats each entry into an arm as a discrete two-choice discrimination (L vs R). We then consider all truns over a 60-min period in terms of overlapping series of four choices (tetragrams; *e.g*., RRRR, RRRL, RRLR etc) (Cleal *et al*., 2021). Data is then split into two formats for analysis: 1) total percentage use (calculated as a proportion of total turns) of each tetragram sequence for the 60 min of exploration – ‘global’ search strategy; 2) search pattern configurations over 10 min time bins throughout the trial, equating to six equal, consecutive time bins – ‘immediate’ search strategy. ‘Global’ search strategy was used to measure of working memory as repetition of previous turn choices must be remembered for patterns of movement to be repeated over 1 h of exploration. The second type of strategy was used to assess cognitive flexibility. For example, animals that adapt their behavioral response to new information would be expected to change behavioral patterns over time. Thus, to assess ‘global’ strategies, number of alternations (RLRL + LRLR) and repetitions (RRR + LLLL) were used as proportion of total number of turns which are highly expressed through 1-hour (% of total turns) (Cleal *et al*., 2021). Average turns were used to assess animals’ locomotion and exploratory activity. The Zantiks AD system (Zantiks Ltd., Cambridge, UK) was used to measure the fish choice for each arm across 1-hour.

### 2.7. Pavlovian fear conditioning

Pavlovian fear conditioning is a behavioral task used to investigate learning and memory in which animal organisms learn to predict aversive event and fear-related memory (Maren, 2001). Zebrafish (*n=* 48) were individually placed in one of four lanes of a tank (25 cm length x 15 cm, 1 L of water) for ~60 min (Valente *et al*., 2012; Brock *et al*., 2017). Fish were habituated for 50-min, in a half check and half grey base screen tank (position switched every 5-min). Baseline preference was recorded for an additional 10-min and assessed by the time spent in the tank areas. Baseline was followed by a conditioning phase in which a conditioned stimulus (CS+; full screen of “check” or “grey”, randomized between each batch) was presented for 1.5-s and followed by a brief mild shock (9 V DC, 80ms; unconditioned stimulus (US)). Subsequently, an 8.5-s of inter-trial interval (ITI) of the non-CS (CS-) exemplar was presented at the bottom of the tank. The CS+/US was repeated eight times. Finally, avoidance of CS+ was assessed by presenting the baseline screen (CS+ and CS–simultaneously) for 1-min, and switching positions after 30-s. The retention index was calculated by the following formula: *retention index = (baseline — probe)* – 1.

### 2.8. Statistics

All data generated in this study are provided online (https://github.com/BarbaraDFontana/ELS_Transgenerational_Zebrafish). Data from the Y-maze protocol was obtained as number of entries into each arm (1, 2, 3 and middle section 4) across a 1-hour trial. To analyze the data according to left and right turns in 10-minute time bins, raw data was processed using the Zantiks Y-maze Analysis Script created specifically for this purpose (available from: https://github.com/thejamesclay/ZANTIKS_YMaze_Analysis_Script).

Normality of data and homogeneity of variance were analyzed by Kolmogorov–Smirnov and Bartlett’s tests, respectively. Because data was normally distributed, one-way analysis of covariance (ANOVA) with CUELS as independent variable (three levels – control (0 days) *vs*. 7 days *vs*. 14 days of CUELS) was used to analyze FMP Y-maze total turns, repetitions, and alternations. Similarly, ANOVA was used to look at results found for novel tank diving task, light-dark preference, Pavlovian fear conditioning and shoal behavior. Tukey’s test was used as post-hoc analysis, data was represented as mean and error of the mean (SEM), and results were considered significant when p ≤ 0.05.

## Results

### CUELS induces changes in anxiety-like behaviors across three generations

There were clear transgenerational effects of CUELS on anxiety-like behaviour. However, this was somewhat nuanced, with 7-days’ exposure (in F_0_) causing differences in all generations, but not 14 days’ exposure. These differences were evident from two separate anxiety tasks (light/dark preference and novel tank test) carried out in two independent batches of fish, suggesting a strong generalizability of the observed effects across context. First, for reference, we show data for the F_0_ fish that was published previously (Fontana et al., 2021), in which there was a clear reduction in anxiety-like behaviour in both the novel tank test, and the light-dark test. **Figure 1** shows the behavioral effects of CUELS for both 7- and 14-days’ exposure, across generations (with F_0_ for reference), in two anxiety-related tasks. There were no effects on distance travelled for any of the exposed animals vs control, suggesting no differences in gross motor activity. However, as was observed in F_0_, F_1_ originated from fish exposed CUELS spent more time spent in the top zone of the novel tank test (ANOVA: F (_2, 44_) = 9.194, *p\*\*\** < 0.001), but this was only the case whether they were exposed to 7 days of CUELS (Tukey test: F_1_ 7 day CUELS vs control: *p\** = 0.0128), not 14 days (Tukey test: F_1_ 14 day CUELS vs control: *p* = 0.5153). F_1_ fish showed similar patterns in the light/dark test, spending less time in the dark (ANOVA: F (_2, 45_) = 5.130, *p\*\** = 0.0098), but this was only significant for F_1_ fish whose parents were exposed to 7 days of CUELS (Tukey test: F_1_ 7 day CUELS vs control: *p\** = 0.0155; Tukey test: F_1_ 14 day CUELS vs control: *p* = 0.9581).

**Figure 1.**
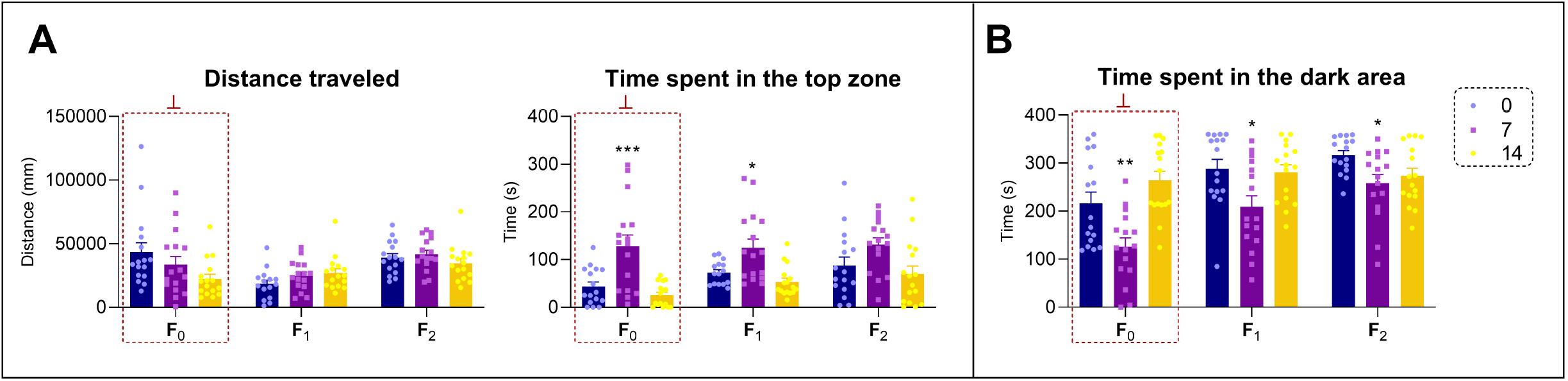
Display the results of ELS in adult zebrafish across multiple generations in two anxiety-related tests: **(A)** Novel Tank Diving Test and **(B)** Light-Dark Test. A significant increase in the time spent in the top area as well as a decrease in time spent in the dark area was observed for the first-generation offspring of 7 days of CUELS (*n* = 15 - 16). The anxiety-like responses observed in the diving tank were attenuated for F_2_ but the effect was still present depending on task (*n* = 16). Asterisks represent significant statistical difference compared to the generation control (0 days of CUELS) (*p\**<0.05,*p\*\**<0.01,*p\*\*\**<0.001). Data were represented as mean ± S.E.M and analyzed by one-way ANOVA for each of the generations, followed by Tukey’s test multiple comparison test. Data in the red box (┴) representing the F_0_ generation have been published elsewhere (Fontana et al., 2021b), and are summarized here for comparison.

The F_2_ generation, as was the case in their parents and grandparents, showed no differences in terms of the distance travelled. Also as with the other generations, CUELS had an impact on F_2_’s anxiety. The F_2_ generation spent less time in the dark compartment (ANOVA: F (_2,45_) = 4.119, *p\** = 0.0228) but only when F_0_ had been exposed to 7 days of CUELS (Tukey test: F_2_ 7 day CUELS vs control: *p\** = 0.0220; Tukey test: F_2_ 14 day CUELS vs control: *p* = 0.7541). In terms of the novel tank test, the F_2_ progeny of CUELS F_0_ spent more time in top (ANOVA: F (_2,45_) = 3.812, *p* =* 0.0295), but the effect was not as clear as in the F_0_ and F_1_ generations. The F_2_ generation that were exposed to 7 days CUELS spent more time in the top only when compared to F_2_ 14 days CUELS (Tukey test: F_2_ 7 day CUELS vs F_0_ 14 day CUELS: *p\** = 0.0267), but not to controls (Tukey test: F_2_ 7 day CUELS vs control: *p* = 0.1523).

### CUELS does not have effects on memory and cognition, fear conditioning or social behavior across generations

As well as anxiety, we also examined several additional cognitive endpoints to examine whether the CUELS had impacts across several domains. Data regarding the F_0_ behavior was previously published (Fontana et al.,2021) and again, are included for reference. No significant differences were observed in F_0_ for social behavior, fear-related memory or working memory in the shoal test, Pavlovian fear conditioning and FMP Y-maze, respectively. Here, we found no effects in any generation from F_0_ animals exposed to CUELS in two memory-related tasks and social behavior. Briefly, for F_1_ no differences in working memory (ANOVA for alternations: F (_2, 45_) = 0.1372, *p* = 0.8722), or repetitive behavior (ANOVA: F (_2, 45_) = 0.2484, *p* = 0.7811, **Figure 2A**), fear conditioning (ANOVA: F (_2, 45_) = 0.370, *p* = 0.6887; **Figure 2B**), or social behavior (ANOVA for shoal area: F (_2, 20_) = 0.6123, *p* = 0.5515; ANOVA for distance between fish: F (_2, 20_) = 0.8135, *p* = 0.4568; **Figure 2C**). Similarly, F_2_ did not show any significant alterations in terms of working memory (ANOVA for alternations: F (_2, 45_) = 0.008, *p* = 0.9920), repetitive behavior (ANOVA: F (_2, 45_) = 0.900, *p* = 0.4138; **Figure 2A**), fear-related memory (ANOVA: F (_2, 45_) = 0.1731, *p* = 0.8416; **Figure 2B**), nor social behavior (ANOVA for shoal area: F (_2, 21_) = 0.2205, *p* = 0.8039; ANOVA for distance between fish: F (_2, 21_) = 0.1063, *p* = 0.8996; **Figure 2C**).

**Figure 2.**
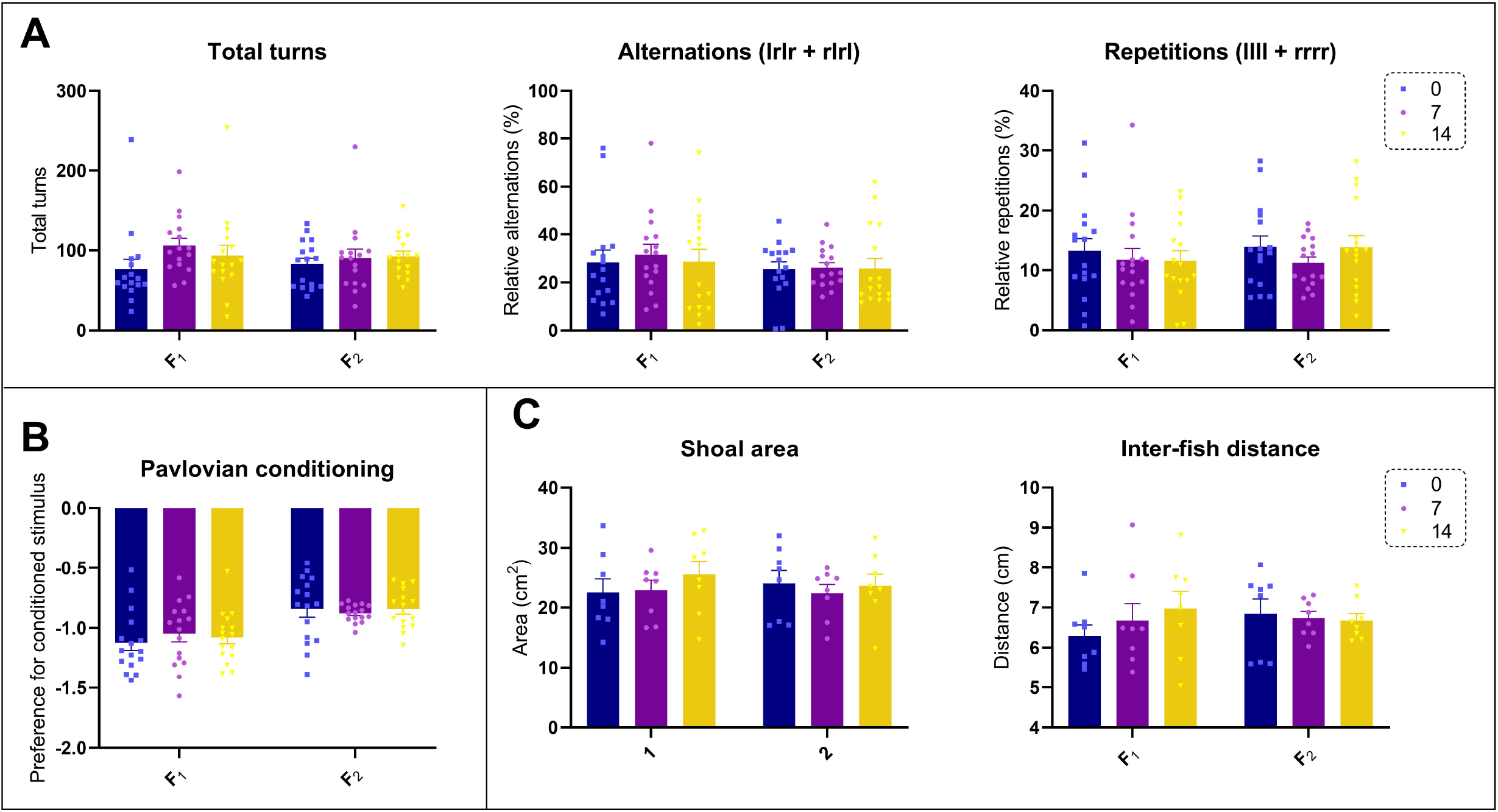
The transgenerational effects of early-life stress (ELS) in (**A**) working memory (*n =* 16), (**B**) fear-related memory (*n =* 16) and (**C**) social behavior (*n =* 8 shoals) in zebrafish. No significant differences were found for all the parameters analyzed. Data were represented as mean ± S.E.M and analyzed by one-way ANOVA.

## Discussion

Here we show, for the first time in the species, that early-life adversity leads to transgenerational inheritance of adapted anxiety-like behavior, to both F_1_ and F_2_ generations, in zebrafish. F_2_ zebrafish bred from F_0_ subjected to 7 days of ELS showed a decrease in anxiety, but only in a single behavioral domain (light-dark preference test, not novel tank test). This may indicate a diminishing, or more domain-specific, effect of ELS on F_2_ (Burggren, 2015). Stressful events may also differentially affect behavior dependent on generation, where different phenotypes are only observed later the consecutive generations (Sagi-Schwartz *et al*., 2008). Based on this, although we have previously seen no evidence of CUELS affecting either social (shoal) behavior or memory and cognition (FMP Y-maze and Pavlovian fear conditioning), these behavioral endpoints were also accessed across-generation. Similarly, no evidence of altered working memory, fear-related memory and social behavior was observed for F_1_ or F_2_, confirming that ELS does not induce any behavioral changes in those behavioral domains that could have skipped generations.

We have previously demonstrated lasting adaptations to stress reactivity following the CUELS protocol (Fontana *et al*., 2021a), where adults exposed as larvae to 7 or 14 days of CUELS showed lower anxiety in a novel environment and in the light/dark test and blunted physiological and behavioral responses to a potent chemical stressor (fear pheromone). In a wild/free-ranging environment, this adaptation may serve to make the fish more proactive when faced with threats (Fontana et al., 2021). However, in the laboratory environment, batch-level variability in behavioral traits may have significant connotations for the reliability of behavioral performance, meaning that within-and between laboratory variability could be impacted, even if the animals themselves were not directly exposed to stressors. This is particularly critical given the high level of in-breeding in most facilities(Tsang *et al*., 2020). Considering the importance of ELS in adult zebrafish behavior, one question remains: does ELS affecting zebrafish adult behavior lead to changes in the germline?

ELS and early-adversity leading to long-last effects in brain and behavior is well established across a range of taxa (Spinelli *et al*., 2009; Schmauss *et al*., 2014; de Vries *et al*., 2018). In rodents, a common model used to investigate the effects of early adversity is maternal separation. This protocol alters neurodevelopmental processes and is a risk factor for the development of psychiatric symptoms later in life (Uchida *et al*., 2010; Schmauss *et al*., 2014; Cui *et al*., 2020; Nishi, 2020). Maternal separation-induced ELS has an impact across generations, affecting DNA methylation in the germline, indicating epigentic modifications are the mechanism of transgenerational transmission (Franklin *et al*., 2010).

Owing to their conserved DNA methylation machinery and mechanisms (Goll & Halpern, 2011), zebrafish have previously proved useful in studying the transgenerational effect of drugs in the environment (Liu *et al*., 2011). The study of the effects of one common endocrine-disrupting contaminant, bisphenol A(BPA), have improved our understanding of the interactions of behavior and epigenetics in zebrafish. BPA disrupted swim activity of larval zebrafish, and also affected two genes (*dnmt1* and *cbs*) which are related to DNA methylation (Olsvik *et al*., 2019). BPA has also been show to cause heritable abnormalities in reproductive tissue, with unexposed F_1_ showing similar differences to exposed F_0_ (Akhter *et al*., 2018). These data suggest that a range of behavioral and physiological impacts of exposure to contaminants can pass across generations, and that epigenetic mechanisms may be underlying it. As well as exposure to environmental contaminents, zebrafish rearing environment has also been shown to affect behavior across generations: F_0_ zebrafish social isolated immediately following fertilization showed decreased aggression compared to group housed fish, and this was then passed to F_1_ (Tamilselvan & Sloman, 2017). Here, we have found that altered anxiety-like behaviors observed in F_0_ fish exposed to ELS appears to be passed into the germline.

Overall, these findings are the first to demonstrate that early-life stress in zebrafish can persistently affect anxiety-like behavior across generations. Importantly, because the differences observed in F_0_ were observed in F_2_, this strongly implicates epigenetic alterations: F_2_ differences must have involved germline-dependent transmission, as F_2_ were never directly or indirectly exposed to ELS (or directly to the F_0_ or F_1_ fish). As such, our findings significantly extend previous data showing the importance of the growing environment, and experienced adversity, in the behavior of future generations. We argue that this should serve as a warning to the zebrafish community that it is a priority to standardize early life environments to try to mitigate. In the future, it would be interesting to examine in more detail the precise epigenetic mechanisms associated with the heritability following ELS in zebrafish. This could potentially help to better understand the mechanisms underlying persistent changes in anxiety induced by early adversity. Zebrafish are a cost-effective alternative to mammals and coupled with their high rates of fecundity, and external fertilization and development of embryos may offer a refined model organism for the study of the long-term effects of ELS.

## Data and Code Availability

The data collected for this study is available at GitHub (github.com/BarbaraDFontana/ELS_Transgenerational_Zebrafish).

## Acknowledgments

This study was supported in part by the CAPES (Brazil) - Finance Code 001 at the University of Portsmouth, UK (BDF). MOP also currently receives funding from the Alzheimer’s Research UK, INTERREG (EU), Dstl (UK), and NC3Rs (UK). The funders had no role in study design, data collection, and analysis, decision to publish, or preparation of the manuscript.

